# The connexin 43 regulator Rotigaptide reduces cytokine-induced cell death in human islets

**DOI:** 10.1101/327171

**Authors:** Seyed Mojtaba Ghiasi, Jakob Bondo Hansen, Dan Ploug Christensen, Thomas Mandrup-Poulsen

## Abstract

**Background:** Intercellular communication mediated by cationic fluxes through the Connexin-family of gap-junctions regulates glucose-stimulated insulin-secretion and beta-cell defense against inflammatory stress. Rotigaptide (RG, ZP123) is a peptide analog that increases intercellular conductance in cardiac muscle-cells by prevention of dephosphorylation and thereby uncoupling of Connexin-43 (Cx43), possibly via action on unidentified protein phosphatases. For this reason, it is being studied in human arrhythmias. It is unknown if RG protects beta-cell function and viability against inflammatory or metabolic stress, a question of considerable translational interest for the treatment of beta-cell failure in diabetes.

**Methods:** Apoptosis was measured in human islets known to express Cx43, treated with RG or the control peptide ZP119 and exposed to glucolipotoxicity or IL-1b + IFNg. INS-1 cells shown to lack Cx43 were used to verify if RG protected human islet-cells via Cx43-coupling. To study mechanisms of action of Cx43-independent effects of RG, NO, IkBa degradation, mitochondrial activity, ROS and insulin mRNA levels were determined.

**Results:** RG reduced cytokine-induced apoptosis ~40% in human islets. In Cx43-deficient INS-1 cells this protective effect was markedly blunted as expected, but unexpectedly RG still modestly reduced apoptosis, and improved mitochondrial function, insulin-2 gene levels and accumulated insulin release. RG reduced NO production in Cx43-deficient INS-1 cells associated with reduced iNOS-expression, suggesting that RG blunts cytokine-induced NF-kB signaling in insulin-producing cells in a Cx43-independent manner.

**Conclusion:** RG reduces cytokine-induced cell-death in human islets. The protective action in Cx43-deficient INS-1 cells suggests a novel inhibitory mechanism of action of RG on NF-kB signaling.

## Highlights

- **Rotigaptide (RG) counteracts cytokine-induced apoptosis in human islets**
- **RG reduces cytokine-induced apoptosis in INS-1 cells independently of Cx43**

## Introduction

By inducing beta-cell endoplasmic reticulum stress and apoptosis via the intrinsic mitochondrial pathway, proinflammatory cytokines have been implicated as mediators of beta-cell failure and destruction causing type 1 diabetes (T1D) and type 2 diabetes (T2D) (1). Antagonism of the action of the prototypic proinflammatory cytokine interleukin-1 (IL-1) improves beta-cell function in T2D patients (2), an effect sustained 39 weeks beyond cessation of the antagonism in responders (3). Despite a strong preclinical rationale (4) anti-IL-1 monotherapy was ineffective overall in recent-onset T1D. However, IL-1 antagonism did moderate inflammation and caused a 2,5-fold higher secretory function in T1D patients with intermediary beta-cell function at baseline (5, 6). Since IL-1 induced beta-cell apoptosis is potentiated by other proinflammatory cytokines such as TNF, IFNg and IL-6, and since anti-TNF therapy improved beta-cell function in a small placebo controlled trial (7), it is likely that a combination of treatments targeting various aspects of signaling caused by the cytokine network is needed to improve efficacy of anti-cytokine strategies in T1D, as has indeed been demonstrated in animal models (8). Thus, there is need for novel safe therapeutic approaches for such combination therapies.

Appropriate pulsatile insulin secretion depends on islet intercellular communication and synchronization (9). Accordingly, cytokine-mediated de-synchronization of intercellular oscillating calcium fluxes alters the beta cell transcriptome and sensitizes it to stress-induced impaired secretory function and apoptosis (10). Gap junctions are intercellular channels composed of two connexin (Cx) hemi-channels, each comprised of six Cx subunits (11). Cx36 is a predominant Cx in beta cells and regulates insulin secretion by calcium flux synchronization (12, 13). Other Cxs are also expressed in islets (Cx43 and Cx45 in mice; Cx30.3, Cx31, Cx31.1, Cx31.9, Cx43, and Cx45 in humans), although the function of these Cxs is less characterized (11, 14, 15). Of note, several studies have failed to show basal Cx43 expression in rat islets or insulin-producing cell lines (16–18).

Whole-body Cx36 deficient mice develop beta-cell destruction and hyperglycemia, whereas beta-cell specific transgenic Cx36 over-expression protects against single high-dose streptozotocin-induced diabetes and restores islet insulin contents in this model. In addition, proinflammatory cytokines reduce beta-cell Cx36 expression, and Cx36 deficiency aggravates cytokine-induced beta-cell toxicity (10). Further, beta-cell specific knockout of Cx43, but not of Cx36, reduces pancreatic insulin content and islet size (19). Therefore, pharmacological targeting of gap junctions has been proposed as a novel approach to rescue pancreatic beta cells from stress-induced apoptosis. An important knowledge gap to guide clinical trials is the demonstration of the importance of the Cx family in human islet apoptosis induced by inflammatory stress.

The peptide Rotigaptide (RG) increases intercellular conductance in cardiac muscle cells and restores gap junction intercellular communication (GJIC) in atrial cardiomyocytes of metabolically stressed rats (20). RG prevents dephosphorylation and thereby uncoupling of Cx43, possibly by acting on yet unidentified protein phosphatases (21, 22). In addition, RG dose-dependently increases the Cx43 expression level in rat cardiomyocytes (23). Here we had two aims: 1) to investigate if Cx43 plays a role in inflammatory-stress induced *human* islet cell apoptosis using RG as a Cx43 coupler 2) to verify that RG exerted an anti-apoptotic effect via Cx43 in human islets by demonstrating loss-of-function of RG on cytokine-induced apoptosis in Cx43 deficient INS-1 cells.

## Methods

### Reagents

Recombinant rat (rr) IL-1b, mouse (rm) IFNg, human (rh) IFNg, and rhTNFa were purchased from R&D Systems (Minneapolis, USA). Rotigaptide (RG; ZP123) and control scrambled inert hexapeptide (CP; ZP119) were provided by Zealand Pharma.

### Cell culture and exposures

The rat INS-1 cell line (generously provided by Claes Wollheim, University of Geneva, Switzerland) known not be express Cx43 (16, 17) was tested negative for *Mycoplasma* and maintained as previously described (24, 25). INS-1 cells were seeded in 6-well plates (1×10^6^ cells/well for RNA isolation), 48-well plates (50,000 cells /well for cell death assay in duplicate) or 96-well plates (30,000 cells/well for MTT assay and ROS assay) (all plates from NUNC, Roskilde, Denmark). After 48 hours of pre-incubation, cells were treated for the time periods indicated with or without 100nM Rotigaptide or CP, for one hour and then cultured with or without cytokine mixture (Cyt) at the concentrations indicated in figures or figure legends, or in glucolipotoxic conditions (0.5uM Palmitate conjugated with 0.1% albumin as described in (26) + 25mM glucose; GLT) for 24 hours. Two concentrations of 100 and 500 nM Rotigaptide were tested in INS-1 cells. No differences in efficacy on cell death and NO well noted and therefore selected 100 nM, in agreement with (23).

### Human islet culture and exposures

Islets from five human heart-beating organ-donors (> 80 purity, donor characteristics listed in table 1) were isolated by the European Consortium for Islet Transplantation (ECIT) in Milan, Italy under local approval and received in fully anonymous form, were cultured in 10% Fetal Bovine Serum (FBS), 1% penicillin/streptomycin (P/S) and 5.6mM glucose at 5% CO_2_ and 37°C as described previously (27). There were no apparent differences in results obtained with islets from male or female donors, and data were therefore combined.

Fifty human islets were pre-cultured in RPMI supplemented with 2% human serum (Life Technologies, Naerum, Denmark), 1% penicillin/streptomycin (P/S) and 5.6mM glucose for 24 hours, treated with or without 500nM Rotigaptide or control peptide for one hour and then exposed to cytokine mixture (300pg/ml rrIL-1b+10ng/ml rhIFNg+10ng/ml rhTNFa) or control medium for 4 days.

**Table 1.** Characteristics of human donors of pancreatic islets.

| No. | Lab Code | Gender | Age | BMI |
| --- | --- | --- | --- | --- |
| 1 | HE-5-15 | F | 22 | 22.8 |
| 2 | HE-10-16 | F | 60 | 23.6 |
| 3 | HE-15-16 | M | 34 | 23.1 |
| 4 | HE-17-16 | M | 50 | 24.8 |
| 5 | HE-2-17 | M | 52 | 25.1 |

### Apoptosis and cell viability assays

For apoptosis assay carried out in duplicate independent cultures, DNA/histone complexes released from the nucleus to the cytosol were measured using Roche cell death assay kit (Roche, Mannheim, Germany) according to the manufacturer’s protocol. As a surrogate of cell viability, mitochondrial function was measured in duplicate by MTT assay in which water soluble MTT (3-(4,5-dimethylthiazol-2-yl)-2,5-diphenyltetrazolium-bromide) is converted by normal cells to an insoluble formazan salt with an optical density (OD) read at 570nm.

### Nitric oxide (NO) assay

As a surrogate of nitric oxide production, accumulated nitrite was measured in duplicate samples from two independent parallel cultures. Supernatants (100ul) from the wells used for INS-1 cell apoptosis assay were mixed with an equal volume of the Griess reagent (one part 0.1% naphtylethylene diamine dihydrochloride and one part 1% sulfanilamide in 5% H3PO4 (Merck, Darmstadt, Germany), and read at 550nm in a plate reader (Thermo Scientific, Naerum, Denmark). The nitrite concentration was calculated using a standard curve of 0.5–40uM concentrations of NaNO_2_ (Merck) as described (28).

### Reactive Oxygen Species (ROS) Assay

ROS was measured in triplicate independent parallel cultures by the 5-(and-6)-chloromethyl-20,70-dichlorodihydrofluorescein (H2DCFDA) (Invitrogen, Naerum, Denmark) probe as previously described (29). Fluorescence was read at excitation 495nm/emission 527nm and results shown as delta fluorescence for the time-interval 90-45 min.

### Real-Time Quantitative RT-PCR

Total RNA was extracted using the Nucleo-Spin kit (Macheray-Nagel, Bethlehem, USA) according to the manufacturer’s instructions. Quality and quantity of the extracted RNA was assessed using a NanoDrop-1000 (Thermo Scientific). Five hundred ng total RNA was used for cDNA synthesis with the iScript^™^-cDNA Kit (BioRad, Copenhagen, Denmark). Real-time qPCR was performed on 12ng cDNA in triplicate with SybrGreen PCR mastermix (Life Technologies) and specific primers (table 2) and run in a Real-Time PCR machine (Applied Biosystems, Naerum, Denmark). Gene expression level was normalized to HPRT1 through -ΔCt analysis.

**Table 2.**
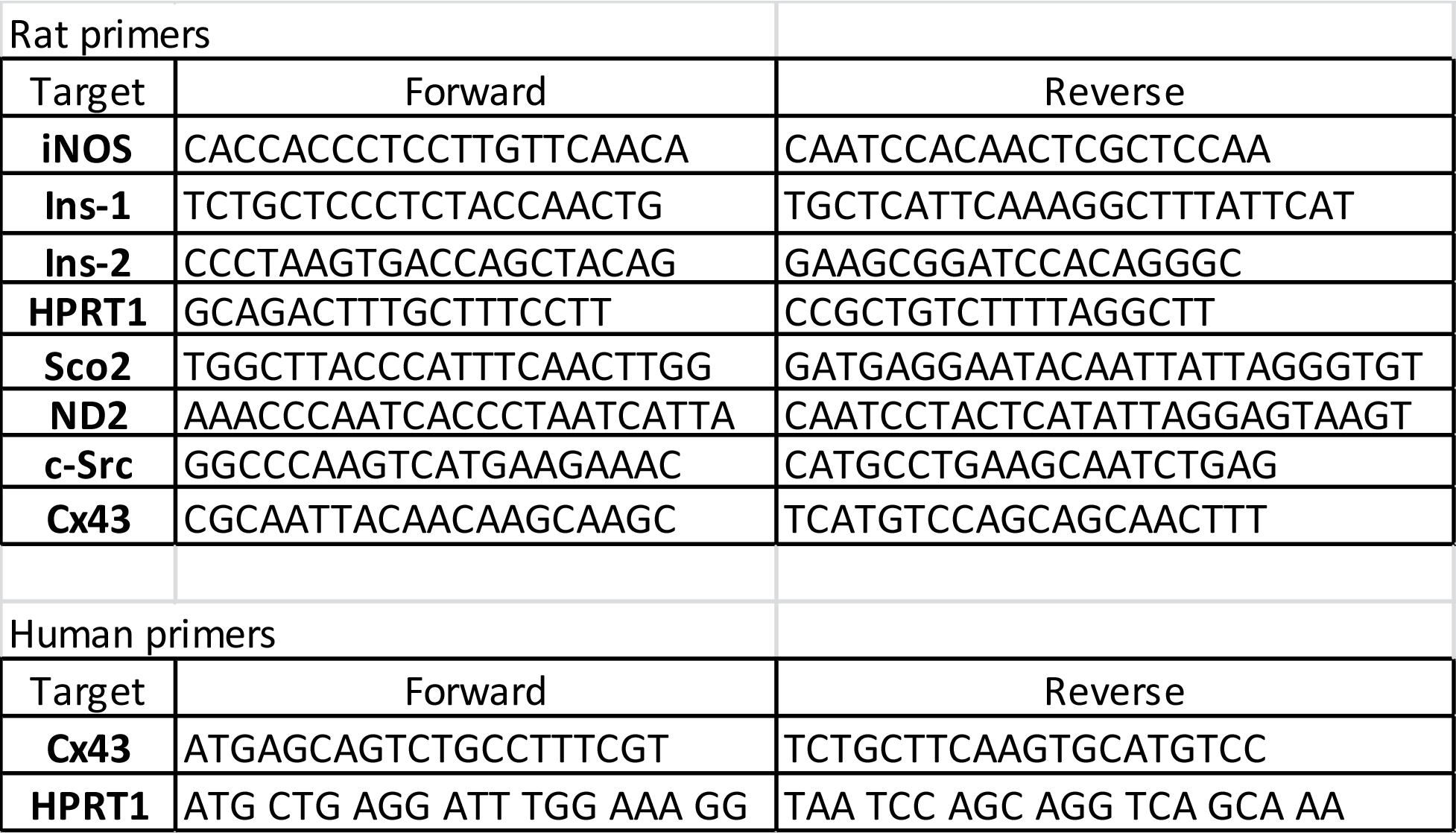
Rat primer sequences.

### Western blot analysis

Cells were lysed on ice with NP-40 lysis buffer containing protease inhibitor cocktail (Life Technologies) and stored at -20 freezer. Lysates were adjusted for protein concentration with Bradford assay according to the manufacturer’s protocol (BioRad), and 50ug protein separated by 4–20% SDS-PAGE, and blotted on PDVF membrane (BioRad). The membranes were stained with anti-IkB (Santa Cruz, Heidelberg, Germany) and alpha-tubulin (Sigma, Copenhagen, Denmark) antibodies and developed with the chemiluminescence detection system Super Signal (Life Technologies) as previously described (30). Light emission was captured using an Alphaimager system (Alpha Innotech, MultiImage III, Broager, Denmark). Band density was quantified using ImageJ software.

### Insulin Assay

Supernatants (1:200 dilution) collected from the INS-1 cell apoptosis assay experiments were used for measurement of accumulated insulin using insulin competitive ELISA assay in duplicate samples from two independent parallel cultures as described (31), except that the enzyme substrate 1-step Ultra TMB (3,3’, 5,5;-tetramethylbenzidine) (Life Technologies) was used here.

### Statistical analysis

Data are presented as means ± SEM, and comparisons between different groups were carried out by ANOVA analysis, followed by Student’s paired *t* test using the GraphPad Prism version 6 (La Jolla, USA). Bonferroni-corrected *P-values* ≤ 0.05 were considered as significant and ≤ 0.10 as a trend.

## Results

### Rotigaptide reduces cytokine-induced apoptosis in Cx43 expressing human islets

Since human islets are known to express Cx43 (fig. 1a), we examined whether the Cx43 activator RG protects human islets from inflammatory-induced cell death. As expected, the cytokine combination increased islet apoptosis by two-fold after 4 days of exposure (fig. 1b). Neither RG nor CP in the absence of cytokines caused apoptosis in human islets. Interestingly, RG significantly reduced cytokine-induced islet apoptosis by 40%, whereas the control peptide did not significantly change cytokine-induced apoptosis

**Figure 1.**
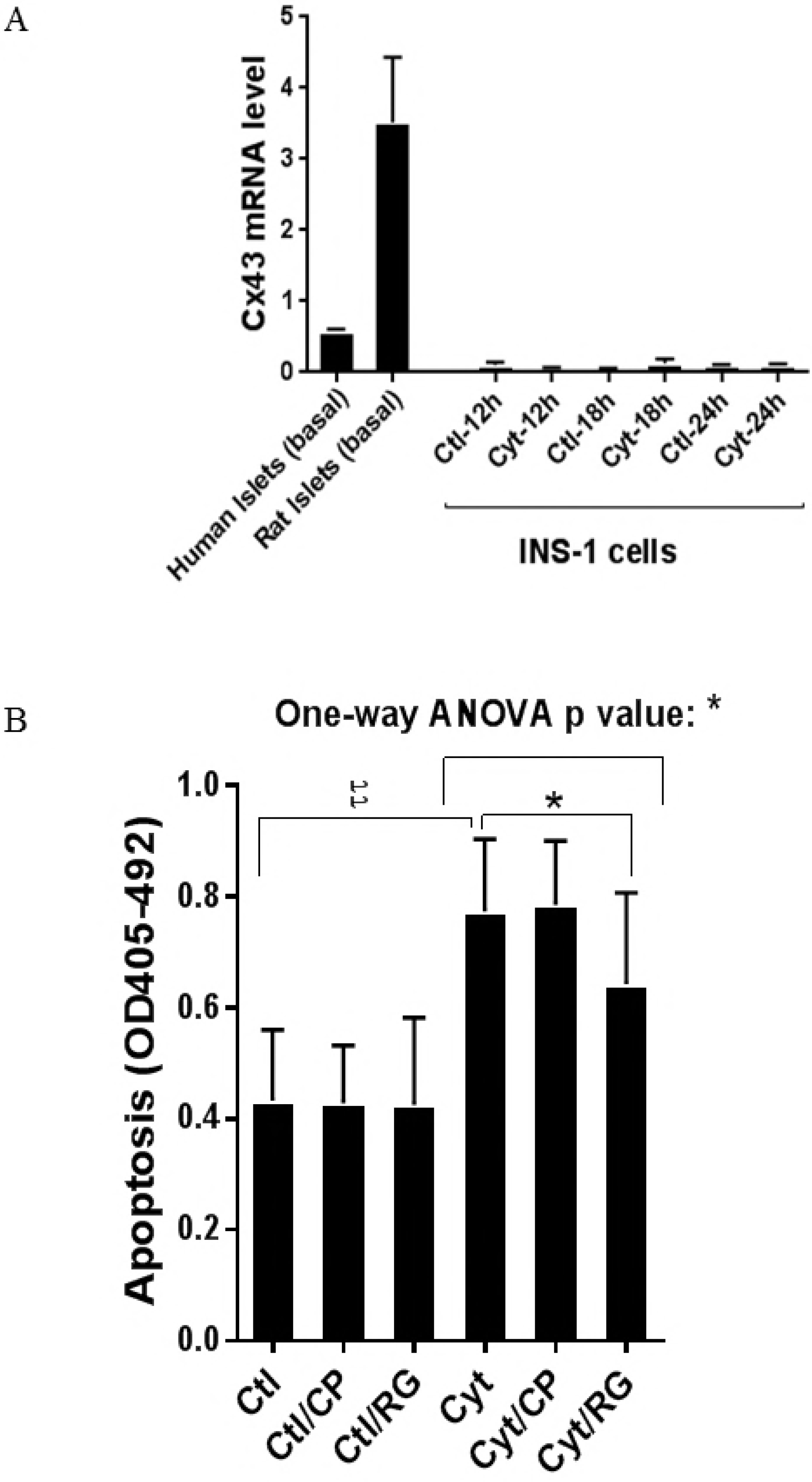
Rotigaptide reduces cytokine-induced apoptosis in Cx43 expressing human islets. Fifty pancreatic islets per condition were pre-incubated with or without 500 nM Rotigaptide/ZP123 (RG) or control peptide ZP119 (CP) for one hour in the presence or absence of cytokine mixture (300 pg/ml IL-1b+10 ng/ml IFNg+10 ng/ml TNFa; Cyt) for 4 days. A) The Cx43 expression level was determined in untreated human and rat islets, and in INS-1 cells exposed to cytokine combination (150 pg/ml IL-1b+0.1 ng/ml IFNg) using specific primers with qPCR. The expression of the genes normalized to HPRT1 was calculated by -ΔCt. B) Apoptosis was measured by Roche cell death assay according to the manufacturer’s protocol. Data are means ± SEM of N=5 donor human islets. * ≤ 0.05, tt ≤ 0.01. The symbols t and star indicate the Bonferroni-corrected paired t-test values of treatments versus control (CTL) and cytokine (Cyt) conditions, respectively. NS: not significant.

### Rotigaptide ameliorates cytokine-induced apoptosis associated with improved mitochondrial function in Cx43 deficient INS-1 cells

In contrast to intact rat islets, the rat insulin-producing INS-1 cells did not express Cx43, and Cx43 was not induced by cytokine exposure for 12–24 hours (fig. 1a). Thus INS-1 cells were used a natural Cx43 loss-of-function cell-model to investigate Cx43 independent effects of rotigaptide on insulin-producing cells. IL-1b concentrations above 15pg/ml combined with a fixed concentration of 0,1 ng/ml IFNg dose-dependently induced apoptosis in INS-1 cells with a peak three-fold induction at 150pg/ml IL-1b (fig. 2a). RG or CP did not by themselves affect INS-1 cell apoptosis. We anticipated that Rotigaptide would not protect Cx43 deficient insulin-producing INS-1 cells against cytokine-induced apoptosis to the same extent as that observed in human islets. Unexpectedly, RG but not CP modestly but significantly reduced apoptosis in cytokine-exposed cells ~10% at IL-1b concentrations above 15pg/ml. Exposure of INS-1 cells to glucolipotoxic conditions significantly increased apoptosis by 3.8 fold, but this was counteracted neither by RG nor CP.

**Figure 2.**
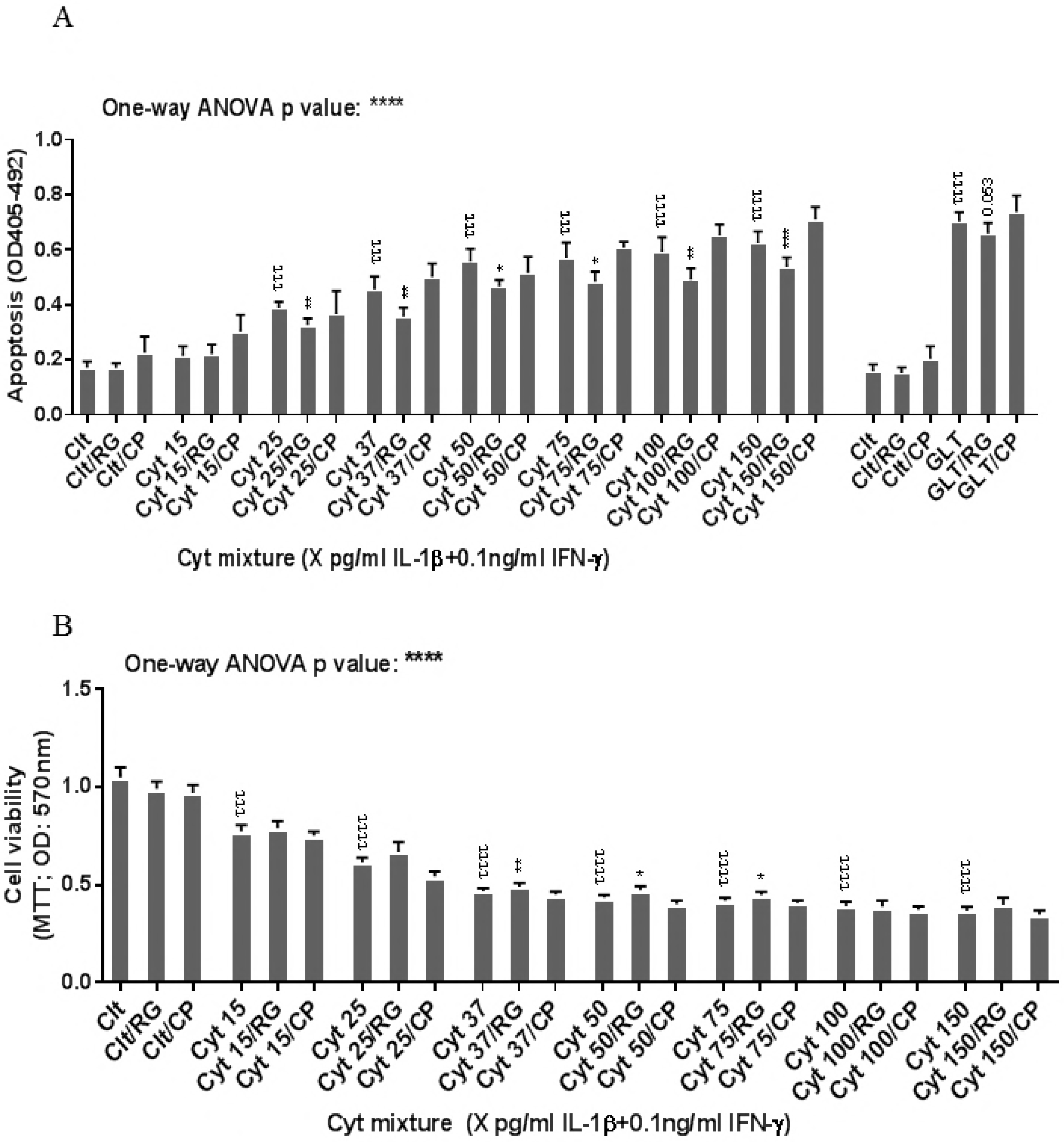
Rotigaptide ameliorates inflammation-induced apoptosis associated with improved mitochondrial function in Cx43 deficient INS-1 cells. INS-1 cells were pre-incubated with or without 100 nM Rotigaptide (RG) or control peptide ZP119 (CP) for one hour in the presence or absence of cytokine combination (IL-1b in the concentrations indicated+0.1ng/ml IFNg; Cyt) or glucolipotoxicity (0.5 uM Palmitate + 25 mM glucose; GLT) for 24 hours. A) Apoptosis was measured by Roche cell death assay according to the manufacturer’s protocol. B) Mitochondrial function was determined using MTT assay. Results are means ± SEM of N=6 independent experiments. * or t ≤ 0.05, ** or tt ≤ 0.01, *** or ttt ≤ 0.001, tttt ≤ 0.0001. The symbols t and star indicate the Bonferroni-corrected paired t-test values of treatments versus control (CTL) and cytokine (Cyt) conditions, respectively.

Since cytokines induce mitochondrial stress in pancreatic beta cells, we then investigated if the Cx43 independent action of RG was associated with prevention of cytokine-induced mitochondrial dysfunction. IL-1b dose-dependently reduced mitochondrial function by 64% at 150pg/ml IL-1b. RG, but not CP, slightly but significantly improved mitochondrial function in cytokine-exposed cells at 37, 50 and 75pg/ml IL-1 (fig. 2b).

### Rotigaptide reduces neither cytokine nor glucolipotoxicity-induced ROS production in INS-1 cells

We next asked if improved mitochondrial function caused by RG was associated with reduced ROS production. IL-1b dose-dependently increased INS-1 cell ROS production (fig. 3a) which was unaffected by RG. Also ROS production in response to glucolipotoxic conditions was unaffected by RG. Since oxidative/nitroxidative stress induced by proinflammatory mediators causes dysfunction of mitochondrial complexes by inhibiting their transcription (32), we explored if RG would prevent cytokine-induced mitochondrial dysfunction. We therefore measured the mRNA levels of two genes encoding key subunits of the mitochondrial complex I and IV: NADH-dehydrogenase subunit 2 (*ND2*) and Cytochrome C Oxidase II (*Sco2*), respectively. Cytokines down-regulated ND2 and there was a trend for down-regulation of Sco2 (p=0.09), but RG did not restore these changes (fig. 3b).

**Figure 3.**
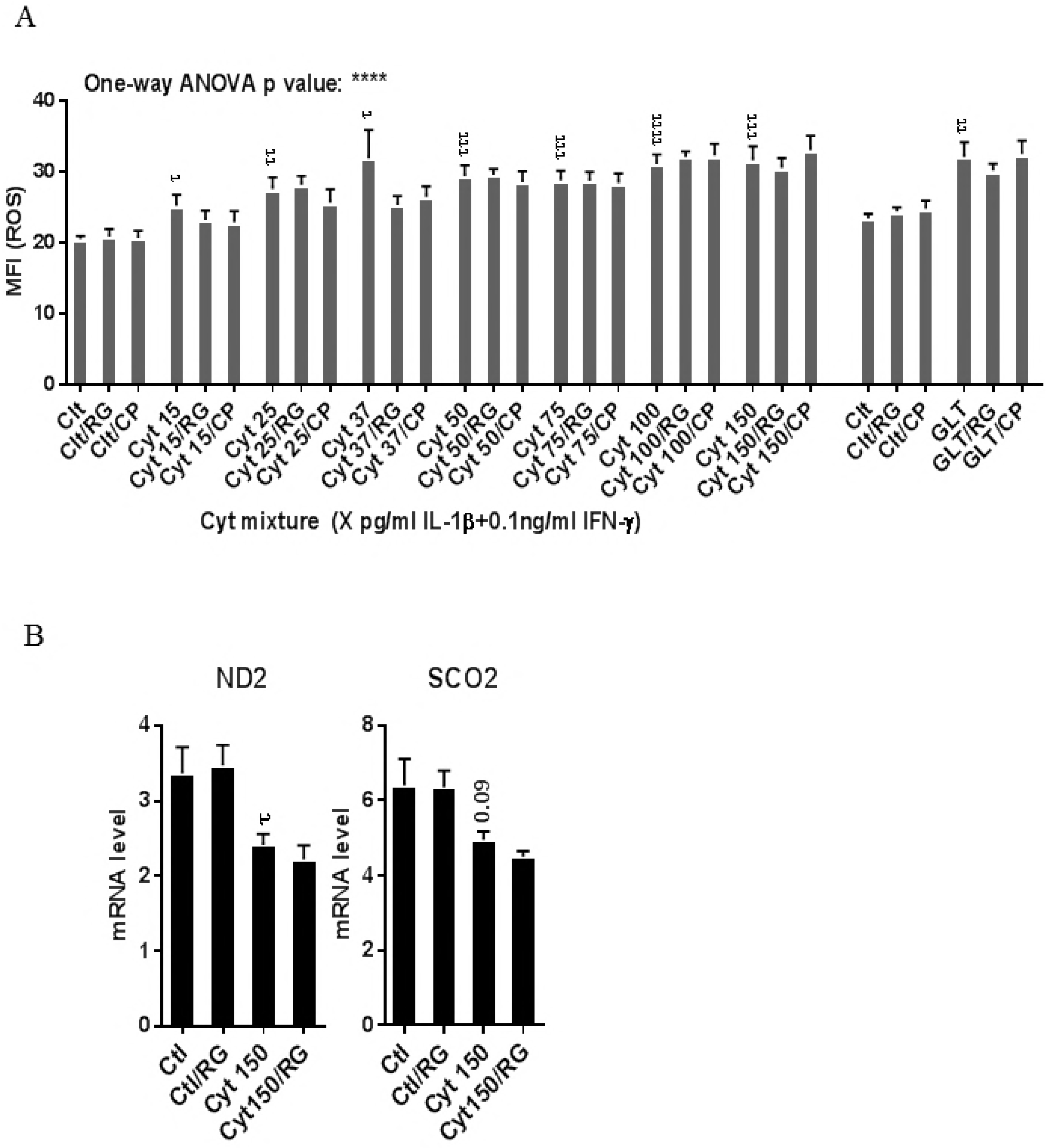
Rotigaptide does not reduce inflammatory or glucolipotoxicity-induced intracellular ROS in INS-1 cells. INS-1 cells were pre-incubated with or without 100 nM Rotigaptide (RG) or control peptide ZP119 (CP) for one hour in the presence or absence of cytokine mixture (IL-1b in the concentrations indicated+0.1ng/ml IFNg; Cyt) or glucolipotoxicity (0.5 uM Palmitate + 25 mM glucose; GLT) for 18 hours. A) Cellular ROS level was determined using DCF assay and presented as MFI. B) The mRNA level of NADH-dehydrogenase subunit 2 (*ND2*) and Cytochrome C Oxidase II (Sco2) genes was determined using specific primers with qPCR. The expression of the genes normalized to HPRT1 was calculated by -ΔCt. Data are the means ± SEM of N=6 (for A)/N=4 (for B) independent experiments. t ≤ 0.05, tt ≤ 0.01, ttt ≤ 0.001, tttt ≤ 0.0001. The symbols t indicates the Bonferroni-corrected paired t-test values of treatments versus control (CTL). ROS: reactive oxygen species, MFI: mean fluorescent intensity, DCF: dichlorodihydrofluorescein.

### Rotigaptide reduces nitroxidative stress independently of Cx43

Next we asked if RG inhibited nitroxidative stress in INS-1 cells. IL-1b significantly induced NO production in INS-1 cells (fig. 4a) which was reduced by RG but not CP at 100 and 150pg/ml IL-1. IL-1b dose-dependently increased iNOS mRNA levels which were reduced by RG but not CP at 100 and 150pg/ml IL-1 (fig. 4b).

**Figure 4.**
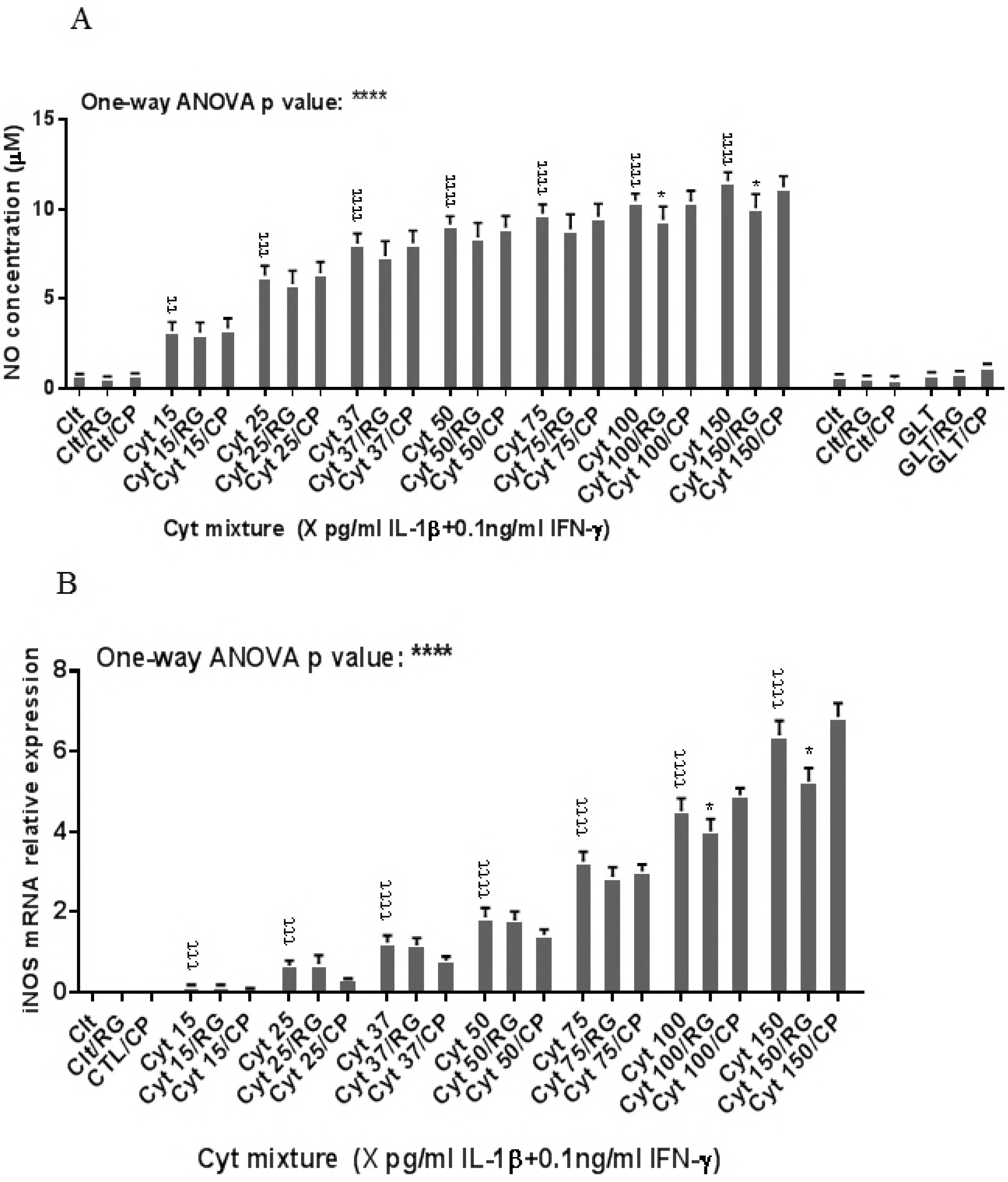
Rotigaptide reduces nitroxidative stress in INS-1 cells. INS-1 cells were pre-incubated with or without 100 nM Rotigaptide (RG) or control peptide ZP119 (CP) for one hour in the presence or absence of cytokine mixture for 24 hours (A), 6 hours (B and D), or in a time course of 5, 10, 15, 20, 25, 30 or 45 min (C). A) Accumulated nitrite was measured with Griess reagent in the supernatant. B and D) iNOS and c-Src mRNA levels were determined by qPCR. The expression of iNOS and c-Src normalized to HPRT1 was calculated with -ΔCt. C) Immunoblot analysis of the time course of cytokine-induced IkB degradation in the presence or absence of RG or CP was quantified with ImageJ software and normalized to tubulin. Data are means ± SEM of N=6 (for A, B and D)/ N=3 (for C) independent experiments. * ≤ 0.05, tt ≤ 0.01, ttt ≤ 0.001, tttt ≤ 0.0001. The symbols t and star indicate the Bonferroni-corrected paired t-test values of treatments versus control (CTL) and cytokine (Cyt) conditions, respectively. AUC: area under curve, NO: nitric oxide, iNOS: inducible nitric oxide synthase, HPRT1: Hypoxanthine-guanine phosphoribosyltransferase 1.

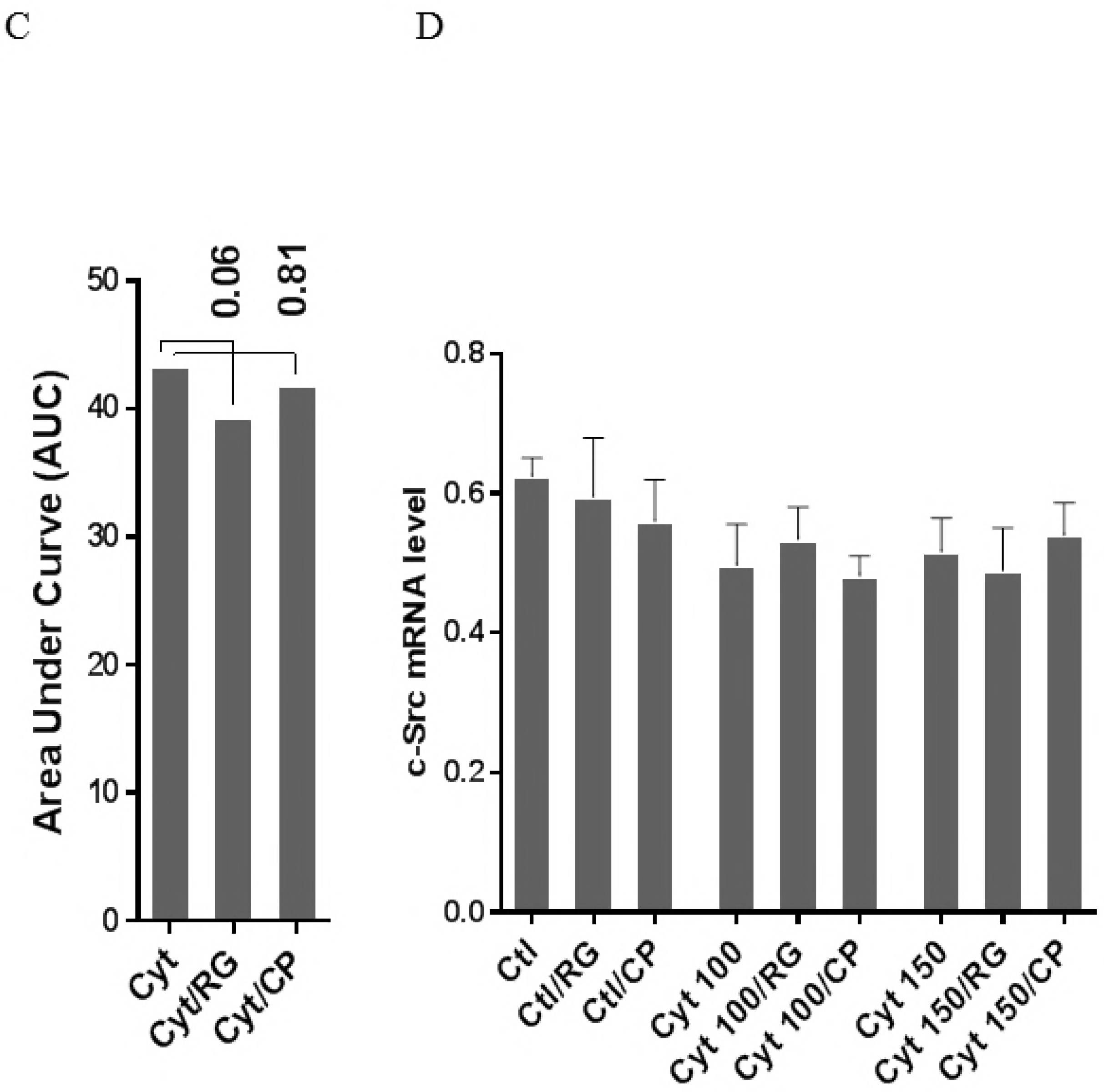

We next investigated if Rotigaptide mitigated nitroxidative stress by impeding IkBa degradation. Cytokines caused IkBa degradation in a time-dependent manner (fig. 4c). There was a strong trend (p=0.06) for RG but not CP diminishing the AUC for IkBa degradation. Since cytokine-induced c-Src levels might be higher in Cx43 deficient cells and since c-Src binds and thereby acts as a sink for IkBa (33, 34), we investigated c-Src expression in INS-1 cells exposed to cytokines in the presence or absence of RG or CP. As shown in fig. 4d c-Src mRNA levels were unaffected by cytokines and RG.

Taken together, the data suggest that Rotigaptide reduces nitroxidative stress independently of its action on Cx43 activity.

### Rotigaptide ameliorates high-concentration cytokine-induced inhibition of insulin secretion and reduction of insulin mRNA in INS-1 cells

To explore if RG restores cytokine-mediated inhibition of insulin biosynthesis and secretion, we measured accumulated insulin secretion and insulin mRNA in INS-1 cells. RG but not CP significantly improved insulin secretion in INS-1 cells at 150pg/ml IL-1b (fig. 5a)

**Figure 5.**
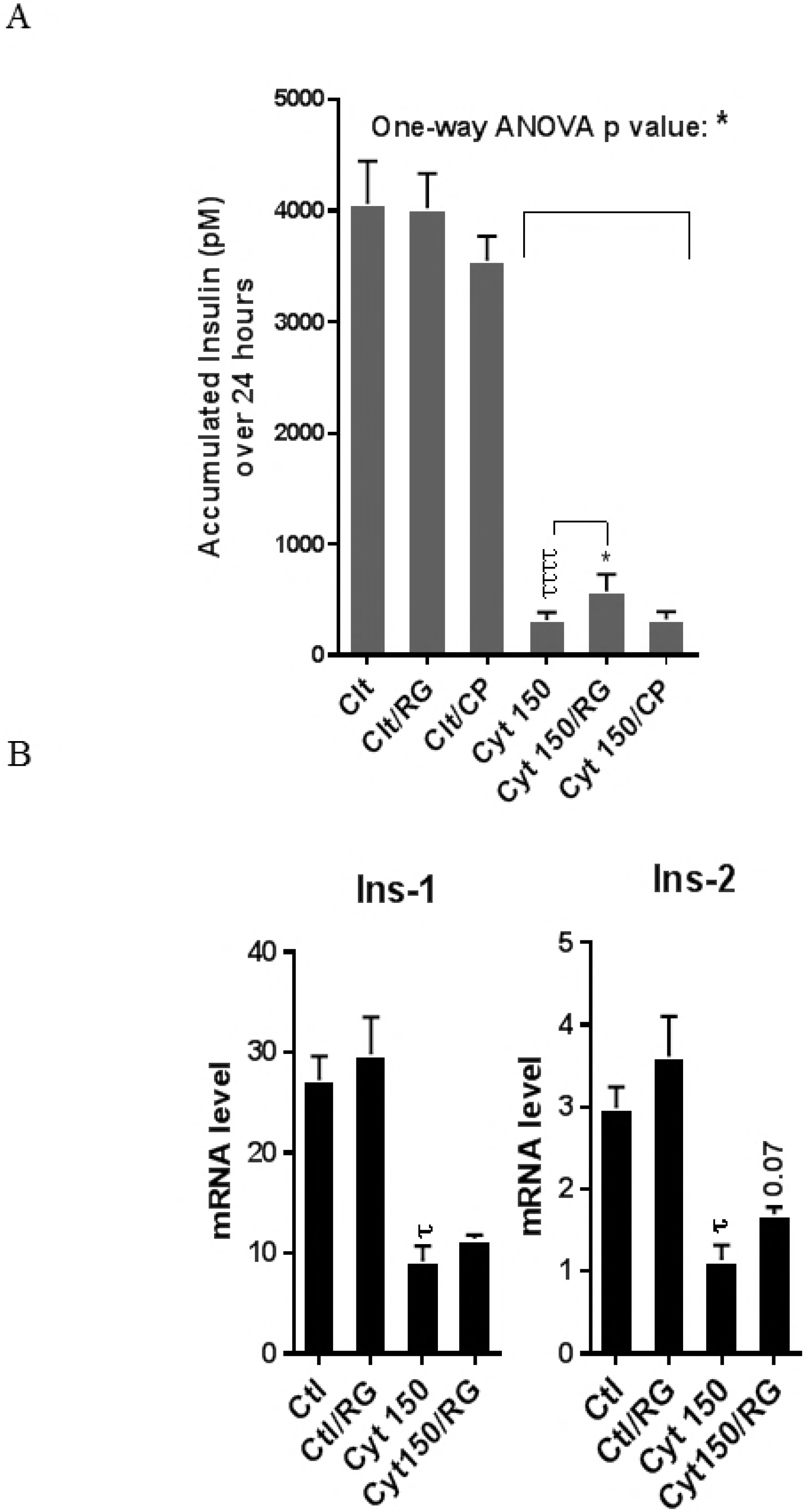
Rotigaptide ameliorates cytokine-induced inhibition of insulin secretion and reduction of insulin mRNA in INS-1 cells. INS-1 cells were pre-incubated with or without 100 nM Rotigaptide (RG) or control peptide ZP119 (CP) for one hour in the presence or absence of cytokine mixture for 24 (A) or 18 hours (B). A) Accumulated insulin was measured by competitive insulin ELISA in the supernatant collected. B) The mRNA level of *insulin-1* (Ins-1), *insulin-2* (Ins-2) genes was determined by qPCR. The expression of the genes normalized to HPRT1 was calculated by -ΔCt. Data are the means ± SEM of N=6 (for A)/ N=4 (for B) independent experiments. * or t ≤ 0.05, **** ≤ 0.0001 (Bonferroni-corrected paired t-test values). t is used for the comparison between CTL and Cyt.

Finally we studied if this partial recovery of secreted insulin was due to increase in the insulin mRNA level. qPCR analysis confirmed significant cytokine-mediated reductions in the mRNA levels of *insulin-1* and *insulin-2*. A trend (p=0.07) for partial reversal of the inhibition in *insulin-2* but not *insulin-1* levels was observed after RG treatment (fig. 5b).

## Discussion

We show here that the Cx43 activator RG reduces proinflammatory cytokine-induced apoptosis in human islets, an observation compatible with the importance of connexins in allowing intercellular cationic fluxes, thereby buffering pro-apoptotic increases in intracellular Ca^2+^ and Fe^2+^ levels mediated by cytokine-induced ER Ca^2+^ release and iron-import (1, 29). Surprisingly, this protective effect was not abrogated in INS-1 cells, known to be Cx43 deficient (fig. 1a) (16, 17). In these cells RG still modestly but significantly reduced apoptosis, and also improved mitochondrial function, insulin-2 gene levels and accumulated insulin release. RG reduced NO production in Cx43 deficient INS-1 cells associated with reduced iNOS expression and IkBa degradation, suggesting that RG blunts cytokine-induced NF-kB signaling in insulin-producing cells in a Cx43 independent manner. High glucose down-regulates Cx43 and inhibits the protein-protein interaction between c-Src and Cx43 in glomerular mesangial cells, thereby promoting binding of c-Src to IkB-alpha, in turn activating the NF-kB pathway (33, 34). These observations raise the intriguing possibility that RG may inhibit NF-kB signaling by enhancing Cx43 expression and activity, but this remains to be shown.

An important question is how RG intercepts NF-kB activation in the absence of Cx43. In Cx43 deficient cells, lack of Cx43/c-Src interaction would be expected to lead to higher c-Src availability for IkBa binding, thereby priming the cells for cytokine-triggered NF-kB activation. We investigated c-Src expression in INS-1 cells and were unable to show altered c-Src mRNA levels caused by either cytokines or RG. In Cx43 competent cells however, this mechanism would be expected only to play a role under conditions of inhibited Cx43 expression, such as those induced by high glucose (33, 34). The lack of protection of RG on high glucose-high lipid toxicity in INS-1 cells may be explained by absence of Cx43, and these experiments should therefore be repeated in human islets, in which RG would be expected to counteract glucose-mediated reduction in Cx43 expression.

Strengths of this study were the use of human islets to demonstrate the protective effect of RG against cytokine-induced apoptosis, and the inclusion of a scrambled peptide as a control for RG. We also took advantage of INS-1 cells known to lack Cx43 expression as a ‘natural’ Cx43 knock-out model (fig.1a) (16, 17), thereby avoiding the risk for transfection artifacts related to shRNA knock-down or CRISPR knock-out.

A limitation was the restricted amounts of human islets available to us for these studies, precluding more mechanistic investigations such as Cx43 activity, Ca^2+^ flux synchronization and studies of glucose-stimulated insulin secretion. Electromobility shift assay, iNOS promotor chromatin immunoprecipitation, or luciferase-based NF-kB reporter assay would help define if NF-kB promoter binding or NF-KB transcriptional activity is reduced in Rotigaptide-treated islets exposed to cytokines. Rotigaptide is also known to increase Cx43 half-life in cardiomyocytes by slowing its trafficking and proteasomal degradation (23, 35, 36). However, proof of Cx43 dependence of such effects of RG in human islets would require knock-down or knock-out of Cx43, which is not effective in intact islets but requires islet dispersion into monolayers, thereby disrupting the normal cell-to-cell contact and communication. Further research is needed to clarify if these effects contribute to the protective action of Rotigaptide in human islets.

When added to the fact that beta-cell specific overexpression of Cx43, but not of Cx36, increases pancreatic insulin content and islet size 19, our observations highlight the translational potential RG as a novel approach to prevent inflammatory beta-cell failure and apoptosis in diabetes and warrant further preclinical studies. The notion that RG may have a dual protective action related to its known activity as a Cx43 activator and to a novel Cx43 independent inhibiting action on NF-kB activation makes RG an attractive candidate for mono- or combination therapy in diabetes.

In conclusion, RG reduces cytokine-induced cell death in human islets, likely by preventing Cx43 uncoupling. RG conferred protection against inflammatory assault even in Cx43 deficient INS-1 cells, suggesting a novel inhibitory mechanism of action of RG on NF-kB signaling. These observations support further development of RG as a novel therapy to protect beta-cell functional mass in diabetes due to its dual protective action on key beta-cell pro-apoptotic pathways.

## Acknowledgments

SMG and TMP were initiators of the study and developed the protocols for the experiments. SMG, JBH and DPC conducted the experiments. All authors discussed data. SMG performed the statistical analysis, constructed figures and tables and wrote the first draft of the manuscript which was edited by TMP, JBH and DPC. This project was funded by Danish Diabetes Academy (DDA), Zealand Pharma A/S and Department of Biomedical Sciences (BMI), University of Copenhagen, the Augustinus Foundation and the Bjarne Jensen Foundation. We appreciate European Consortium for Islet Transplantation (ECIT), Milan, Italy, for providing donor human islets and Zealand Pharma A/S for providing the ZP123 and ZP119 peptides. Rie Schultz Hansen and Adam Steensberg are thanked for helpful advice and information regarding the peptides.

